# Longitudinal analysis of retinal ganglion cell damage at individual axon bundle level in mice using visible-light optical coherence tomography fibergraphy

**DOI:** 10.1101/2022.11.15.516181

**Authors:** Marta Grannonico, David A. Miller, Jingyi Gao, Kara M. McHaney, Mingna Liu, Michael Krause, Peter A. Netland, Hao F. Zhang, Xiaorong Liu

## Abstract

We developed a new analytic tool based on visible-light optical coherence tomography fibergraphy (vis-OCTF) to longitudinally track individual axon bundle transformation following acute optic nerve crush injury (ONC) in mice. We analyzed four parameters: lateral bundle width, axial bundle height, cross-sectional area, and the shape of individual bundles. We showed that axon bundles became wider and thicker at 3-days post ONC. The bundle swelling at 3-days post-ONC has correlated with about 15% retinal ganglion cell (RGC) soma loss. At 6-days post-ONC, axon bundles showed a significant reduction in lateral width and cross-sectional area, followed by a reduction in bundle height at 9-days post-ONC. Bundle shrinking at 9-days post-ONC has correlated with about 68% RGC soma loss. Both experimental and simulated results suggested that the cross-sectional area of individual RGC axon bundles is more sensitive than the bundle width and height to indicate RGC soma loss. This study is the first to track and quantify individual RGC axon bundles *in vivo* following ONC injury and establish the correlation between the morphological changes of RGC axon bundles and RGC soma loss.

## Introduction

Retinal ganglion cell (RGC) loss is a hallmark of optic neuropathies, such as glaucoma [1-3], and neurodegenerative diseases that affect vision, such as Parkinson’s and Alzheimer’s disease [4, 5]. Thus, recognizing RGC loss at its earliest stage is crucial to prevent further irreversible vision loss [6-8]. However, current clinical diagnostic methods to detect the functional and structural changes in the inner retina are often not sensitive or specific enough to directly assess RGC health [9, 10]. An essential parameter for glaucoma diagnosis is visual field (VF) loss; however, studies suggest that more than 30% of RGCs may be lost before detection on VF testing [11-15]. Characteristic changes in the optic nerve head (ONH) and optic disc are also used for glaucoma diagnosis, but identification of damage can be subjective, and grading varies between observers [16]. Among the clinical noninvasive imaging modalities [17-20], optical coherence tomography (OCT) is most widely used for diagnosing and monitoring optic neuropathies. OCT’s cross-sectional imaging capabilities enable measurement of the retinal nerve fiber layer (RNFL) and ganglion cell – inner plexiform layer (GCIPL) thickness *in vivo* as indirect indicators of RGC health [14, 21, 22]. However, inconsistent axial resolution (5 μm to 10 μm) and segmentation algorithm discrepancy among different clinical devices make estimating RGC health less reliable [23, 24]. In conventional RNFL analysis, algorithm failure due to edge smoothing may also cause underestimation of focal areas of thinning when they do occur over time [25].

Importantly, the clinical parameters measured by OCT imaging such as the RNFL or GCIPL thinning are not specific indicators for RGC damage. First, a significant variability in RGC density and RNFL thickness exists among individual patients. For example, the RNFL thickness of healthy human subjects varies from 50 μm to about 120 μm as measured by OCT [26]. Overlaps of RNFL ranges were also observed between healthy subjects and glaucoma patients [26]. Shin and colleagues followed 292 eyes from 192 patients with primary open-angle glaucoma (POAG). They found that only 72 eyes (24.7%) showed progressive GCIPL thinning, and among the 72 eyes, 41 eyes showed progressing visual field (VF) loss [27]. The different patterns of disease progression thus emphasize the need to monitor individual patients longitudinally [22, 27]. Secondly, the ganglion cell layer (GCL) contains non-RGC cells, the displaced amacrine cells (DACs), which vary from 30% to 80% in the GCL; DACs are largely unaffected by glaucoma [2, 28]. We observed 38% RGC axon loss at 3-5 days (d) post optic nerve crush (ONC) injury, but at the same time, we found no significant reduction in overall RNFL thickness in mice [29, 30]. In addition, the inner plexiform layer (IPL) contains synaptic connections among different retinal cell types in rodent and primate eyes [28, 31, 32]. In other words, the RNFL or GCIPL thinning may not be sensitive enough to detect RGC damage specifically. Therefore, there is a need for more accurate and sensitive biomarkers for RGC damages following disease insult.

We recently applied visible-light OCT (vis-OCT), which operates from 510 nm to 610 nm and reaches an axial resolution of 1.3 μm, in the mouse retina [33, 34]. Because optical scattering contrast nearly exponentially decays with increasing wavelength, the visible-light spectral range offers a much higher contrast than the near-infrared (NIR) range [34]. As a result, vis-OCT provides unique anatomical and functional imaging capabilities that facilitate RGC damage evaluation [33]. Recently, we developed vis-OCT fibergraphy (vis-OCTF) to visualize and quantify changes in individual RGC axon bundles in mice [34-36]. In this study, we applied vis-OCTF to track the changes of axon bundles following the ONC injury by quantifying four parameters: (1) lateral bundle width, (2) axial bundle height, (3) cross-sectional area, and (4) bundle shape. We correlated the *in vivo* changes in RGC axon bundle morphology with RGC soma loss after the ONC injury. Based on the experimental data, we created a numerical simulation for each parameter to determine the sensitivity and the floor value - a threshold at which no further change is observable [37, 38].

## Results

### Establishing a new analytic tool for identifying and tracking individual axon bundles *in vivo*

Taking advantage of the improved axial resolution offered by vis-OCTF and associated data processing methods [34], we established an analytic tool for tracking changes in individual RGC axon bundles rather than the bulk thickness of the RNFL *in vivo*. Briefly, four vis-OCT volumes were acquired from the same eye with the optic nerve head (ONH) aligned to each of the four corners of the field of view. This minimized the curvature of the eye and maximized the reflectance of the RNFL [34]. Figure 1A shows one of the four processed vis-OCT fibergrams from an adult wildtype mouse. The red arc shows the path of a resampled circumpapillary scan [35, 36] centered at the ONH with two RGC axon bundles highlighted by 1 and 2 (Fig. 1B). The width of each bundle was measured using the signal intensity profile, as shown in Fig. 1C, which gives 25.0 μm for bundle 1 and 11.7 μm for bundle 2. The resampled circumpapillary B-scan reconstructed at 425 μm radius from the ONH shows the cross-sectional view of axon bundles 1 and 2, as indicated by the dashed green lines. The axial bundle height was measured using the normalized intensity peaks, which give 19.7 μm for bundle 1 and 12.8 μm for bundle 2, respectively (Fig. 1D).

**Figure 1.**
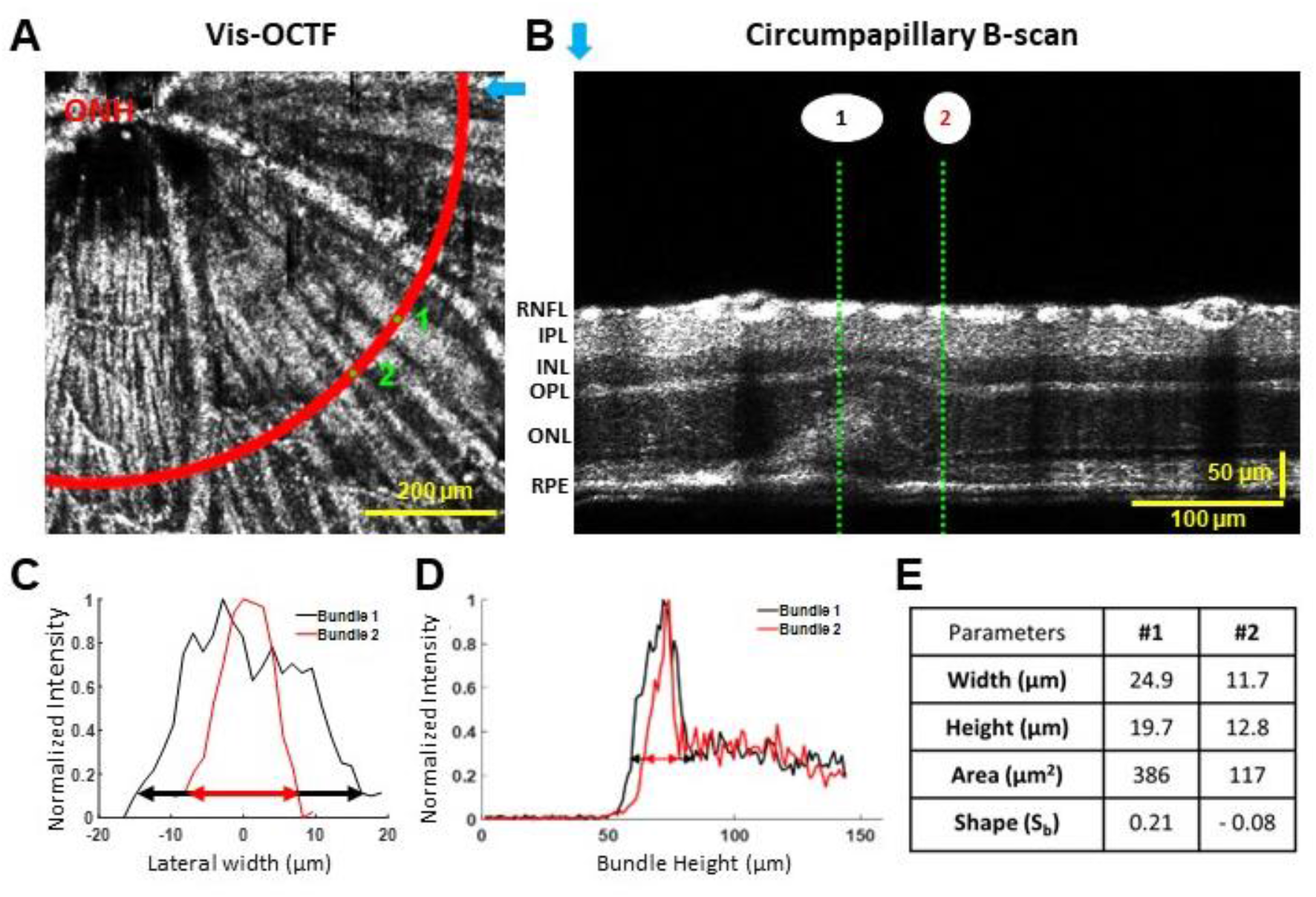
*In vivo* identification and quantification of RGC axon bundle morphology. (A) A fibergram from a single OCT volume of a wildtype mouse. Two RGC axon bundles (1) and (2) were labeled at the radius of 425 μm from the ONH as indicated by the red arc. (B) Circumpapillary B-scan image reconstructed along the red arc in A shows the cross-sectional image of the retina. The blue arrow in A indicates the leftmost A-line in B. The green dashed lines indicate the bundles (1) and (2). (C-D) Intensity profile of the lateral width (C) and the axial intensity (D) of the bundles (1) and (2). The width of the axon bundle is measured at 1/e^2^ decay, as indicated by the black (1) and red (2) arrows. (E) Table of the lateral width, bundle height, cross-sectional area, and the shape indicators of bundles (1) and (2). ONH: optic nerve head; RNFL: retinal nerve fiber layer; IPL: inner plexiform layer; INL: inner nuclear layer; OPL: outer plexiform layer; ONL: outer nuclear layer; RPE: retinal pigment epithelium.

We quantified the cross-sectional area using two methods: the pixel-based cross-sectional area and approximated cross-sectional area from the ellipse (See Methods section for more details). The mean of 57 individual axon bundles measured by pixel area was 158.1 ± 69.1 μm^2^, consistent with mean calculated as ellipse (160.0 ± 82.3 μm^2^, p=0.90, Student’s t-test). We used calculated ellipse area for the rest of the studies (Fig. 1). As summarized in Fig. 1E, the areas of the axon bundles 1 and 2 are 386 μm^2^ and 117 μm^2^, respectively. In addition, we developed a dimensionless indicator for the shape of bundle (S_b_), which normalizes the bundle width to height ratio between -1 and +1 (see Method section for more details). A wider axon bundle, such as bundle 1, has a positive shape value (S_b1_ = 0.21); whereas a thicker axon bundle, such as bundle 2, has a negative shape value (S_b2_ = - 0.08). Altogether, we established four parameters to detect changes *in vivo* in axon bundle morphology: width, height, area, and shape.

### *In vivo* tracking of morphological changes in RGC axon bundles after ONC

We applied vis-OCTF to examine the structural changes at the single axon bundle level following ONC injury. Figure 2A shows example fibergrams and circumpapillary B-scans of a mouse retina acquired at baseline (before ONC), 3-d, 6-d, 9-d, and 15-d post-ONC (pONC). The red arc indicates the path of the resampled circumpapillary B-scan at the radius of 425 μm from the ONH. The blue boxes in the middle panel in Fig. 2A show the magnified views of the fibergrams with five axon bundles tracked over time. The right panels of Fig. 2A show the circumpapillary scans resampled at the same location at different time points. At baseline, the five axon bundles exhibited diverse width, height, area, and shape (Fig. 2B). Following ONC injury, the width of bundle 1 (blue) increased from 23.5 μm (before ONC) to 25.0 μm at 3-d, and then progressively decreased at 6-d pONC (19.9 μm); by contrast, the width of bundle 4 (green) decreased from 23.6 μm (before ONC) to 21.3 μm at 3-d and continue decreased at 6-d pONC (20.4 μm). The height also changed following different patterns. For example, the height of bundle 1 increased from 11.6 μm (before ONC) to 20.3 μm at 3-d and 20.6 μm at 6-d pONC, and then decreased at 9-d pONC. The height of bundle 4 increased from 10.3 μm (before ONC) to 16.8 μm at 3-d pONC and then decreased to 15.7 μm at 6-d pONC. The cross-sectional bundle area presented an overall clear pattern of increase-to-decrease from bundles 1 to 4, although the peak point for each bundle is different. Bundle 5, on the other hand, remained stable until 9-d pONC (Fig. 2B). The shape, however, did not show a clear trend for the selected bundles, which is likely due to different rates of changes in width and height for individual bundles. These observations suggest that (1) axon bundles have diverse morphology; (2) four of the five bundles labeled here exhibit an initial swelling phase before a shrinking phase; and (3) the four parameters of individual bundles change following different patterns.

**Figure 2.**
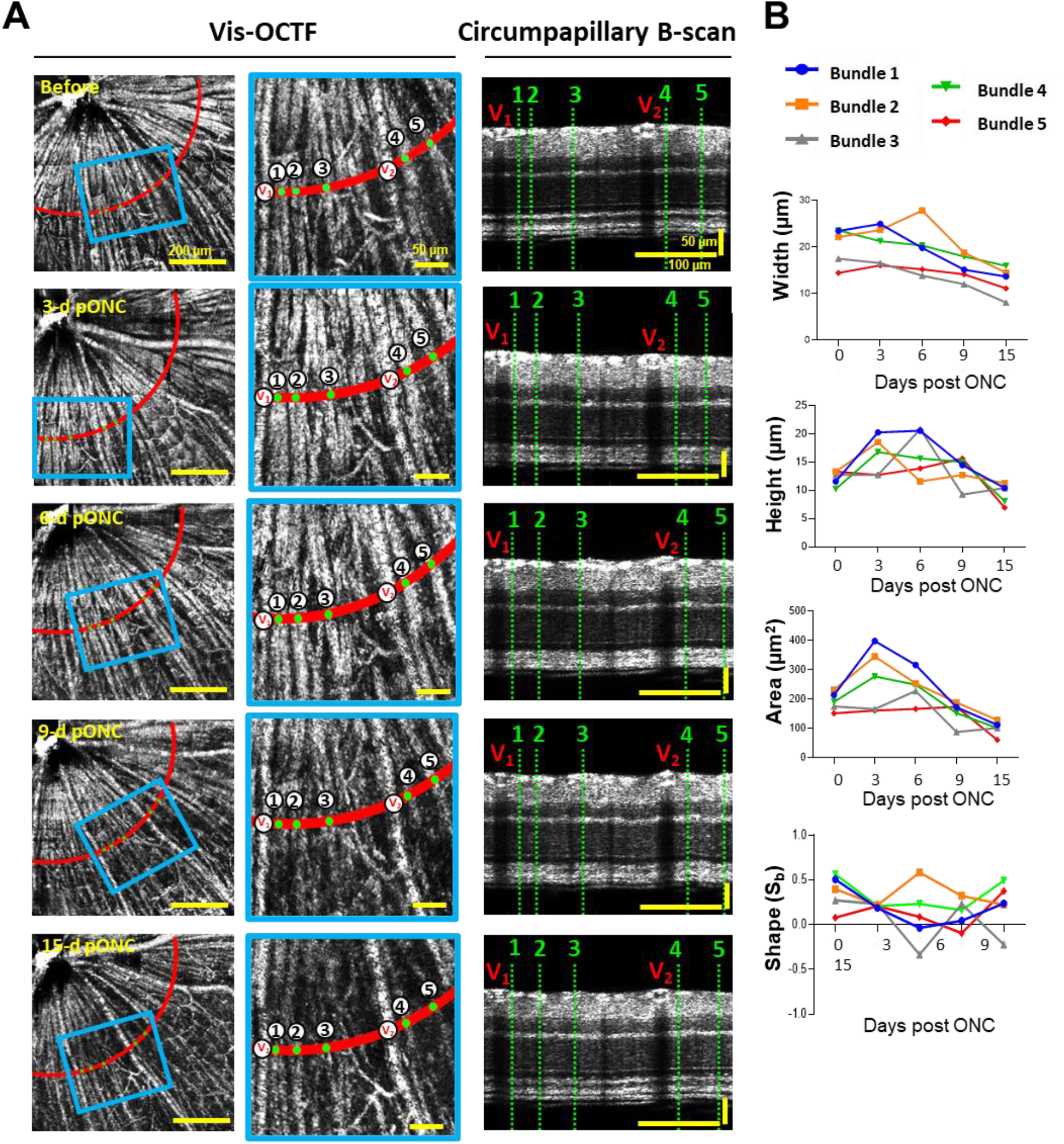
*In vivo* tracking of individual axon bundles following optic nerve crush (ONC) injury. (A) *In vivo* fibergram image of the same retina at baseline and 3-days (d), 6-d, 9-d, and 15-d post ONC (pONC). The left two panels are the vis-OCTF images and right panel is the resampled circumpapillary B-scans at the radius of 425 μm (red arc). Middle panels show the magnified views of highlighted regions in left panels (blue boxes). The green points (1-5) indicate the same five axon bundles tracked over time. The red arrows indicate blood vessels (V_1_ and V_2_). (B) Quantification plots of the width, height, area, and shape (S_b_) of the five tracked axon bundles over time.

We identified and tracked 141 axon bundles from 3 mice following ONC injury. We plotted the histograms of the four parameters (Suppl-Fig. 1) and their smoothed distributions are shown in Figure 3. The RGC axon bundle width was significantly increased from baseline (black, mean: 13.7 ± 4.6 μm) to 3-d pONC (dark red; 14.5 ± 4.5 μm; p = 4.6e-2, One-way ANOVA), followed by a decrease at 6-d pONC (red; 12.8 ± 4.3 μm, p = 5.9e-2) and 9-d pONC (light red; 11.5 ± 3.8 μm, p<1e-4, Fig. 3A). Similarly, the height distribution plot shown in Figure 3B also indicates a significant increase of bundle thickness at 3-d (black; baseline mean: 12.5 ± 3.9 μm; dark blue; 3-d pONC: 13.5 ± 4.5 μm, p=1.5e-2). However, the height returned to baseline at 6-days after ONC (blue; 13.1 ± 4.3 μm, p= 2.5e-1), followed by a significant decrease at 9-d pONC (light blue; 11.1 ± 4.0 μm, p=5.0e-4).

**Figure 3.**
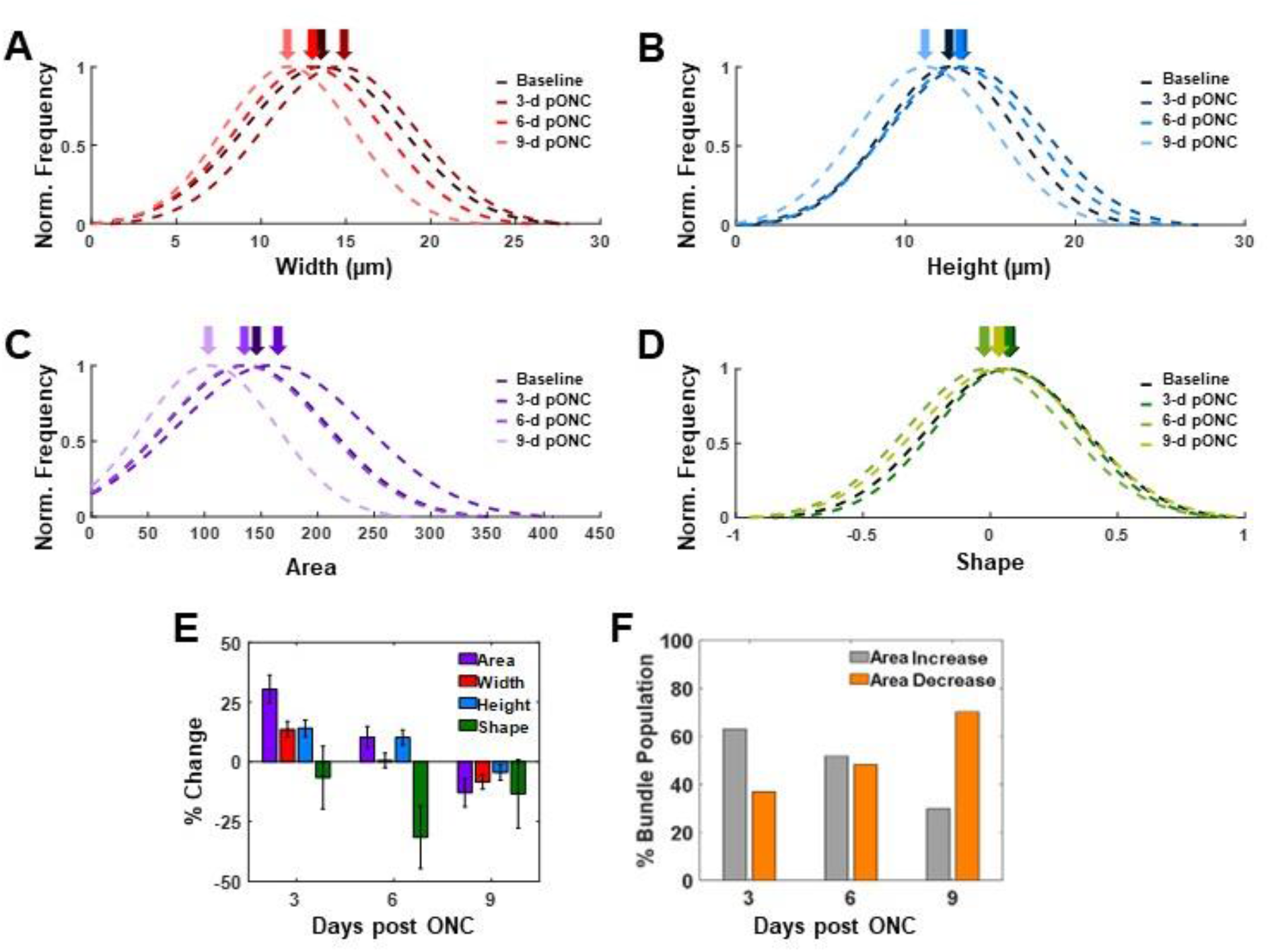
Distribution of the changes in RGC Axon Bundle morphology following the ONC injury. (A-D) Smoothed distributions of the lateral width (A), axial height (B), cross-sectional area (C), and shape (D) for 141 axon bundles (n = 3 mice) tracked over time following the ONC injury (pONC). Shaded arrows point to the mean value of the distribution curve. (E) Percent change of RGC axon bundle width (red), thickness (blue), and area (purple) with respect to baseline values. (F) Percentage of axon bundles exhibiting increased (gray) or decreased (orange) cross-sectional area at different times compared to baseline.

Figure 3C shows the distribution of the bundle area: at 3-d pONC (dark purple), axon bundles presented larger size compared to baseline (black; baseline: 136.6 ± 71.7 μm^2^; 3-d pONC: 158.6 ± 82.9 μm^2^; p = 9.0e-4). At 6-d pONC (purple), the area returned to baseline (6-d pONC: 133.8 ± 71.3 μm^2^; p = 9.1e-1), where at 9-d pONC the area of the axon bundles (light purple) was significantly smaller compared to the baseline area (9-d pONC: 103.8 ± 58.9 μm^2^, p<0.0001). We also noticed that large bundles tend to elongate axially right after the ONC injury (blue arrows in Suppl-Fig. 1). The distribution curves of the axon bundle shape show a progressive shift toward the negative values from baseline (black; 0.065 ± 0.3) to 3-d (dark green; 0.067 ± 0.3, p = 9.9e-1), 6-d (green; 0.011 ± 0.3, p = 2.8e-2), and 9-d pONC (light green; - 0.037 ± 0.3, p = 4.8e-2, Fig. 3D), suggesting that axon bundles naturally have a broader shape that begin to shrink to a more circular shape after injury.

We calculated the percent change of width, height, area, and shape of the 141 tracked axon bundles (Figs. 3E-F). The width of the axon bundles (red) was14% higher at 3-d pONC and only 0.5% higher at 6-d pONC compared to baseline. The height of the axon bundles (blue) was 13% higher at 3-d and 9% higher at 6-d pONC compared to baseline. By 9-d pONC, the width and height of RGC axon bundles decreased by 10% and 7%, respectively. In other words, the width dropped below baseline close to 6-d pONC, while height dropped below baseline between 6-d and 9-d pONC. The cross-sectional bundle area, which enhances the subtle changes in the axon bundle morphology, reveals a clear trend (Fig. 3E). The area increased by 30% at 3-d, and 8% at 6-d, and then decreased by 16% at 9-d pONC, compared to baseline. We also observed a reduction in bundle shape from baseline by 7% at 3-d, then to 32% at 6-d followed by 13% at 9-d (Fig. 3E). Figure 3F shows changes in size in the overall axon bundle population. About 60% of bundle population presented a size increase at 3-d pONC, and 70% of bundle population showed a size decrease at 9-d pONC. At 6-d pONC, about half of bundle population increased (51%) and half of bundle population decreased (48%).

As we have demonstrated that the RNFL thickness measurements were not as sensitive as RGC axon bundle measurements to detect early RGC loss [35, 39, 40], we next compared the GCIPL measurements with our new axon bundle parameters (Fig. 4). In mice, the GCIPL includes RNFL, GCL, and IPL as shown in Figure 4A. We quantified the GCIPL thickness from the circumpapillary B-scan reconstructed at the radius of 425 um (Fig. 4A). The smoothed distribution of the GCIPL measurements (n=120) shows that the GCIPL thickness was not affected at 3-d pONC compared to the baseline (baseline, black, 60.3±5.2 μm; 3-d pONC, dark blue, 61.7±6.1 μm; p= 0.26; Fig. 4C). While at 6-d pONC (blue), and 9-d pONC (light blue), the GCIPL thickness was significantly decreased to 56.0±4.8 μm (p<0.0001) and 51.8±4.5 μm (p<0.0001), respectively, compared to the baseline. In other words, the GCIPL thickness is also not as sensitive as our axon bundle parameters to detect RGC damage.

**Figure 4.**
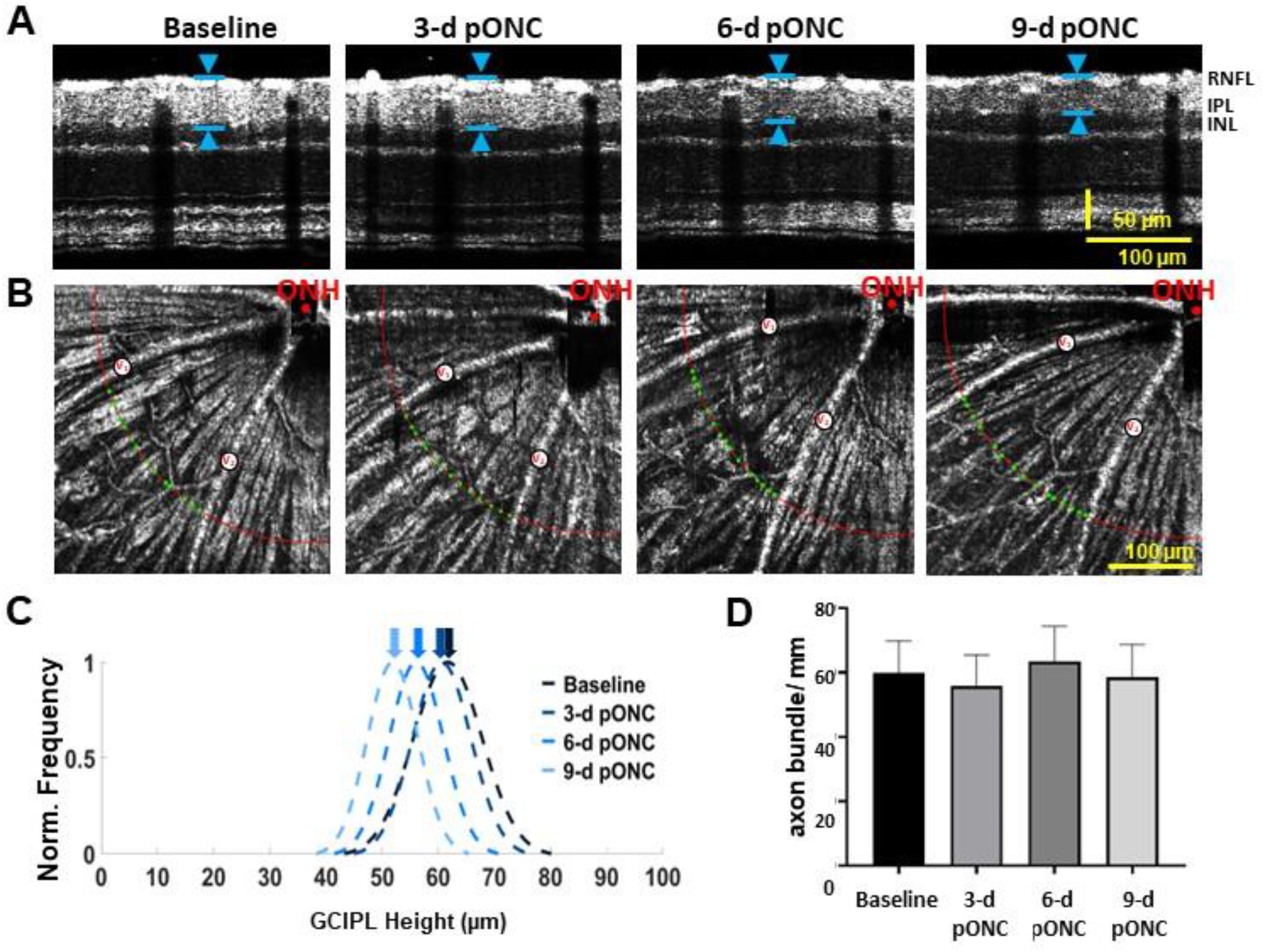
No significant change in the GCIPL thickness was detected at 3-days post ONC by vis-OCT imaging. (A) Resampled circumpapillary B-scans from the same location from at different time points. Blue arrows indicate GCIPL axial thickness. (B) *In vivo* fibergram image of the same retina at baseline and 3-days (d), 6-d, and 9-d post ONC (pONC). V_1_ and V_2_: blood vessels. Green dots indicate axon bundles between V_1_ and V_2_ at the radius of 425 μm (red arc). ONH: optic nerve head. (C) Smoothed distribution of the GCIPL axial height measurements. (D) Average of the number of RGC axon bundle per mm at baseline and 3-days (d), 6-d, and 9-d post ONC.

In addition, we tracked the changes of the number of axon bundles (Fig. 4D). Before ONC, the density of RGC axon bundles was 60 ± 9 axon/mm. It was not changed significantly after ONC (3-d pONC, 56±9 axon/mm; 6-d pONC, 63±10 axon/mm; and 9-d pONC, 58 ± 10 axon/mm; p>0.5, One-way ANOVA). As the overall RGC axon bundle pattern remain largely normal after the injury, our data suggest that the overall RNFL thickness may not change significantly. The conventional measurements of the layer thickness thus may not be able to detect subtle changes in individual axon bundles in optic neuropathies [14, 21, 22].

Taken together, we showed that (1) RGC axon bundle morphology is a more sensitive indicator for RGC damages than the GCIPL thickness; (2) a majority of the axon bundles experience a swollen phase followed by a shrinking phase following ONC injury; (3) the change in width of the axon bundles is more sensitive than the change in the height immediately following ONC injury; (4) the RGC axon bundle cross-sectional area, which combined the width and height measurements, amplified the subtle changes in axon bundle morphology; and (5) the RGC axon bundle shape parameter showed a strong decrease from baseline suggesting a shift towards bundles becoming elongated axially.

### Confocal microscopy imaging confirmed axon bundle damages after ONC

We performed confocal microscopy to validate the axon bundle damages following ONC injury. The retina was dissected and double-immunostained with mouse anti-tubulin beta 3 (Tuj1) and rat anti-neurofilament H (NFH) (Suppl-Fig. 2) [41]. We confirmed that somas of degenerating RGCs became labeled by NFH at 3-d pONC (top panel, Suppl-Fig. 2). Confocal images in Suppl-Fig. 3 were taken at the ONH region close to the lesion site of the ONC surgery. In the control retina, axon bundles were well organized and directly converged toward the ONH (Suppl-Fig. 3A). At 3-d pONC, some axon bundles became disorganized with loose or splitting fibers (yellow arrows, Suppl-Fig. 3B). We also observed retraction bulbs, the non-growing counterparts of growth cones, at the tip of the lesioned axons (white arrows) [42]. At 9-d pONC, the axon bundles near the ONH became entangled, with some retraction bulbs and lesioned axon tips pointing away from the ONH (Suppl-Fig. 3C). At 15-d pONC, the overall bundle structure degenerated at the ONH (Suppl-Fig. 3D).

### Establishing the correlation of morphological changes in axon bundles with RGC soma loss

We next determined the correlation between the *in vivo* vis-OCTF parameters and the RGC soma loss by *ex vivo* confocal imaging. For this set of experiments, we acquired vis-OCT images from healthy wild type mice and mice at 3-d, 9-d, and 15-d pONC, respectively. Immediately following vis-OCT imaging, mice were sacrificed, perfused, and the retinas were dissected and immunostained with rbpms, an RGC marker [43]. Flat-mounted retinas were then imaged by confocal microscopy [35, 43]. As demonstrated in the schematic representation of the flattened retina in Fig. 5A, we divided the retina into superior (S), inferior (I), nasal (N), and temporal (T) quadrants. Examples of *in vivo* vis-OCTFs (left panel) and their corresponding *ex vivo* confocal microscopy images of flattened retinas (middle panel) stained with rbpms are shown side by side in Figs. 5B-E. Magnified views of retinal confocal images show a steady decrease in RGC soma density compared to control at 3-d, 9-d, and 15-d pONC (right panel in Figs. 5B-E).

**Figure 5.**
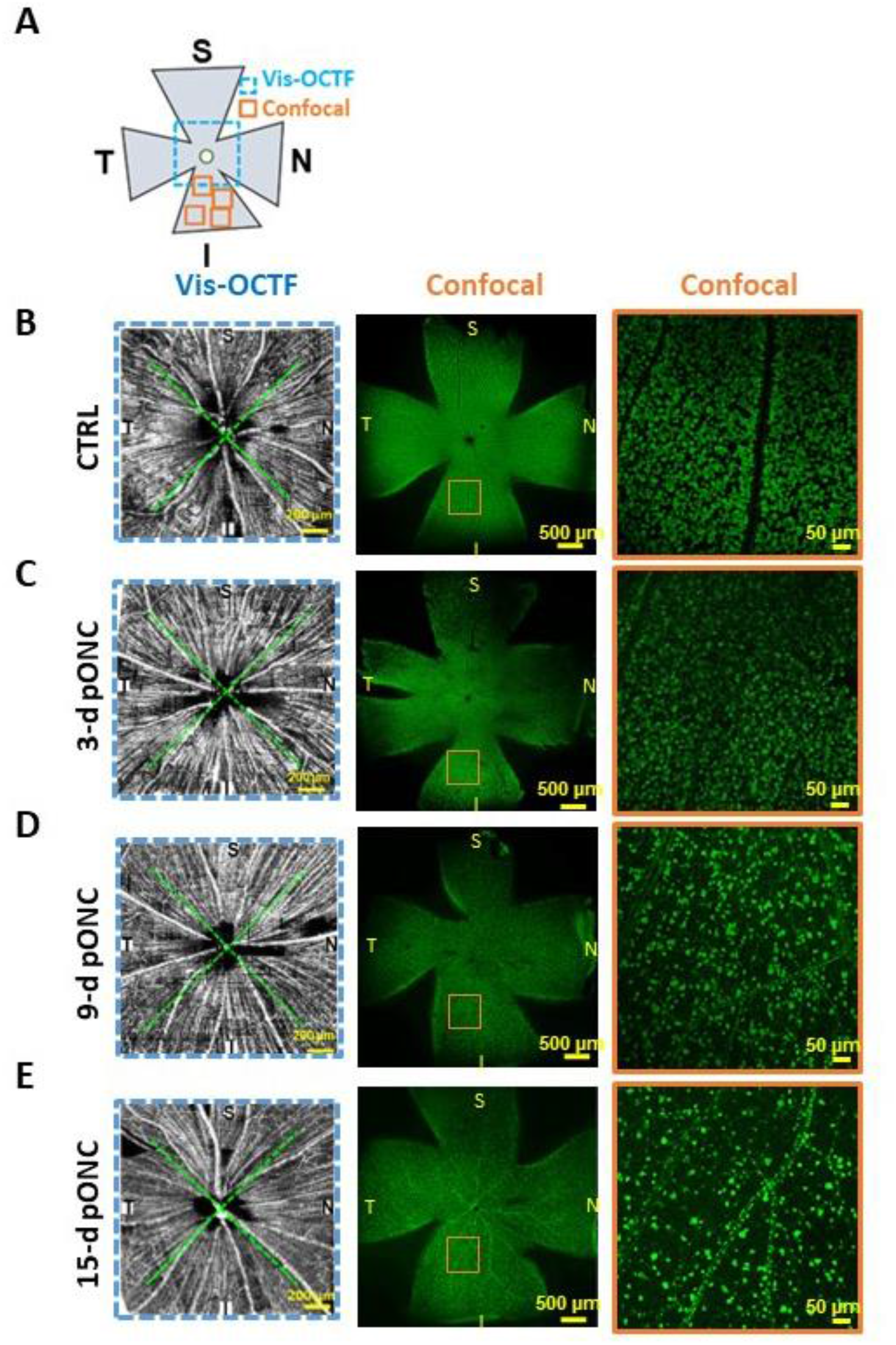
Confocal imaging of flat mounted retinas for quantification of RGC loss following vis-OCT imaging. (A) Schematic representation of the flat mounted retina with vis-OCT (blue box) and confocal microscopy (orange boxes) FOVs overlaid. (B-E) Vis-OCTF (left panel) followed by confocal microscopy images of RGCs labeled by rbpms for estimating RGC density in control (B), 3-d (C), 9-d (D) and 15-d (E) pONC eyes. Right panels show the magnified views of highlighted regions in middle panels (orange boxes). Green dashed lines in vis-OCTF divided the field of view into 4 regions: superior (S), nasal (N), temporal (T) and inferior (I).

We plotted the rbpms-positive RGC density of each retina as a function of each of the four axon bundle parameters quantified from the vis-OCT images (Figs. 6A-D). Each data point represents the average reading per retina. The mean density of rbpms positive RGCs in controls was 4095 ± 209 cells/mm^2^ (n=6 retinas). At 3-d pONC, the mean density was decreased to 3475 ± 343 cells/mm^2^ (n=9 retinas), a 15% reduction on average. The means density of rbpms continued to decrease with time: 1305 ± 104 cells/mm^2^ (68.1% reduction, n=8 retinas) at 9-d pONC, and 611 ± 37 cells/mm^2^ (85.1% reduction, n=3 retinas) at 15-d pONC.

**Figure 6.**
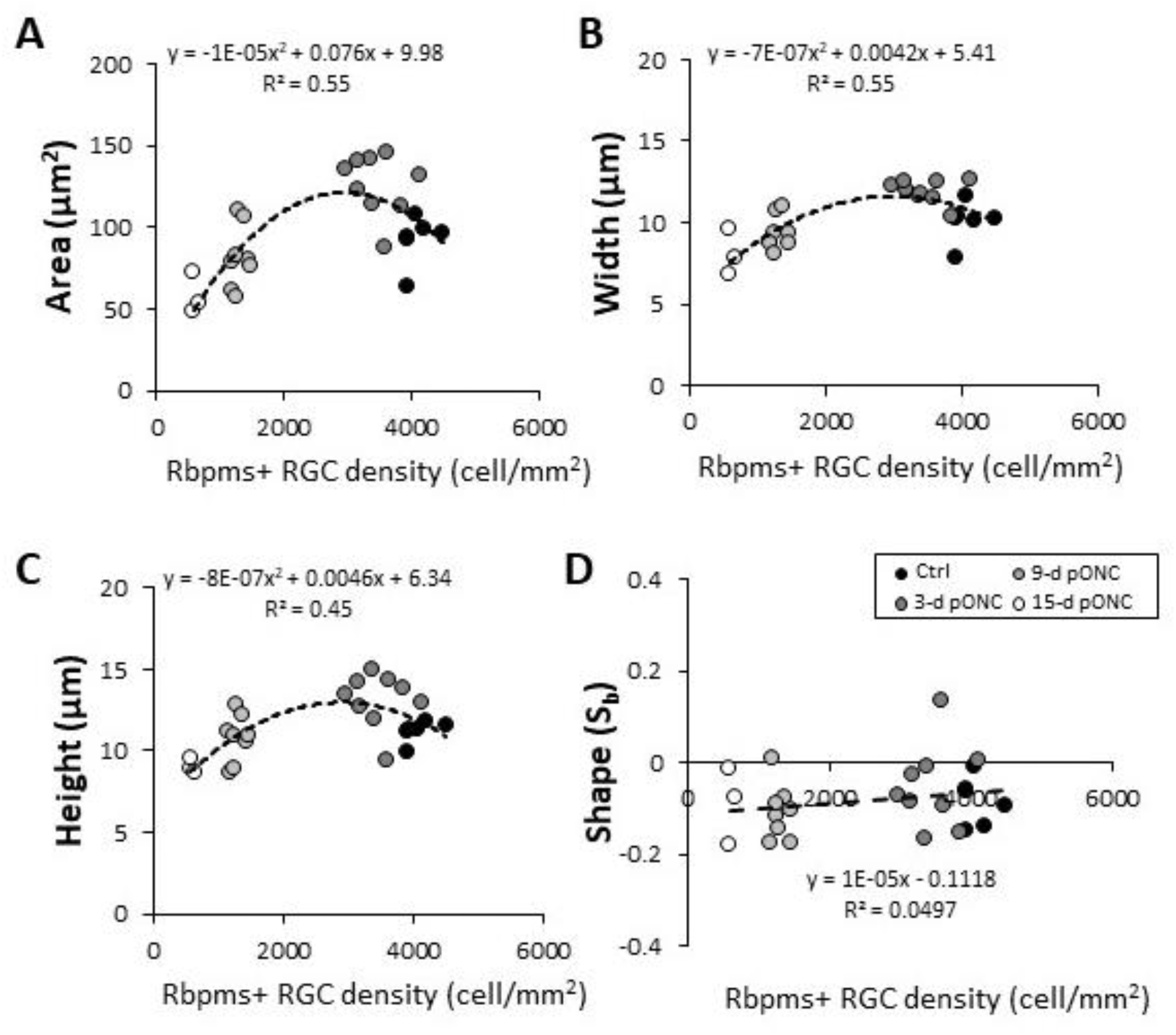
Correlation of RGC soma density and axon bundle size measurements. (A-D) RGC axon bundle cross-sectional area (A), width (B), height (C), and shape (D) plotted as a function of rbpms + RGCs density for each retina. Black, dark grey, light grey, and white dots represented data from control (N=6), 3-d (N=9), 9-d (N=8) and 15-d (N=3) pONC, respectively. Second order polynomial regression (dotted line) were fitted to the data for (A-C), and linear regression were fitter to (D). The equation and r-square values are labeled on the figure.

The overall relationship between axon bundle height, width, and area with RGC density was non-linear (Figs. 6A-C), mainly due to a swelling phase that immediately followed ONC. For example, at 3-d pONC (Fig. 6A), a15% of RGCs lost (p<0.001) correspond to 36% increase of the axon bundles area compared to the control (control: 92.4 ± 13.6 μm^2^, n= 6; 3-d pONC: 126 ± 17.8 μm^2^, n=9; p <0.01). At 9-d pONC, 68% of RGCs lost (p<0.001, compare to the controls), correspond to a 12% reduction in the axon bundle area (81.9 ± 17.8 μm^2^, n=8, p=0.5). At 15-d pONC, the axon bundle area decreased by 37% (58.4 ± 10.4 μm^2^, n=3; p=0.03), while RGC density suffered an 85% reduction (p<0.001, Fig. 6A).

We found that initially the axon bundle’s width, height, and area were negatively correlated with RGC density and around 9-d post ONC morphological changes of axon bundles the and RGC density start becoming positively correlated. To estimate where the positive correlation begins for each parameter, we used the second-order polynomial regression model to fit the bundle width plot (R^2^=0.55, p < 0.001), height plot (R^2^=0.45, p<0.001), and area plot (R^2^=0.55, p<0.001) (dashed lines in Figs. 6A-C; also see Table 1). From the regressions, we found that the axon bundle area peaked at 152 μm^2^ with an RGC density of 3780 cells/mm^2^ (8% cell loss), while the width and height both peaked at a lower RGC density of 3000 cells/mm^2^ (27% cell loss) and 2875 cells/mm^2^ (30% cell loss), respectively, reinforcing the idea that the area might represent a more valuable parameter to track subtle changes in axon bundle morphology after injury. We did not observe a clear trend of shape changes in axon bundle changed after ONC injury (Fig. 6D). The shape parameter remained stable slightly below zero in control and ONC retinas, to which we fit a linear regression model (p=0.47, One-way ANOVA; Fig. 6D). The R-squared values, significances, and estimates of peak values are listed in Table 1. These results suggest that (1) the width, height, and area parameters of the axon bundles change with RGC loss following a similar pattern but with different level of sensitivity; and (2) area is the first parameter to show a positive correlation with RGC loss following ONC injury.

**Table 1.** Regression analysis of RGC axon bundle morphology and soma loss (see Figure 5)

| Parameter | Equation | R<sup>2</sup> | p | Peak of the Curve/ tipping point |  |  |
| --- | --- | --- | --- | --- | --- | --- |
|  |  |  |  | RGC density<br>(cells/mm<sup>2</sup>) | axon<br>parameters | RGC loss<br>(%) |
| **Area** | $y = -1E-05x^2 + 0.076x + 9.98$ | 0.55 | < 0.001 | 3780 | 152.9 $\mu\text{m}^2$ | 7.7 |
| **Width** | $y = -7E-07x^2 + 0.0042x + 5.41$ | 0.55 | < 0.001 | 3000 | 11.8 $\mu\text{m}$ | 26.7 |
| **Height** | $y = -8E-07x^2 + 0.0046x + 6.33$ | 0.45 | < 0.001 | 2875 | 12.9 $\mu\text{m}$ | 27.8 |
| **Shape** | $y = 1E-05x - 0.11$ | 0.05 | 0.27 | N/A | N/A | N/A |

### Location-dependent changes in axon bundle morphology after ONC

Due to the unpredictability of location and the extent of RGC damage caused by the crush surgery, we compared the relationship between bundle morphology and RGC density in different retinal quadrants after ONC. In Figure 7, the four axon bundle parameters for the superior (S), inferior (I), nasal (N), and temporal (T) regions of the retina were plotted as a function of the RGC density for each orientation. Second-order polynomial regression models fitted the data to each region (dashed lines, Figs. 7A-C). The R-square values, significances, and estimates of peak values for all four regions were listed in Table 2 (all regressions were statistically significant, except for temporal width regression). For example, at 3-d pONC, the average density of RGCs had significantly decreased by 20% in the superior leaflet (3314 ± 356 cells/mm^2^ for control, and 2681 ± 337 cells/mm^2^ at 3-d pONC, p<0.01), and significantly decreased by 13% in the nasal leaflet (4371 ± 334 cells/mm^2^ for control, and 3826 ± 385 cells/mm^2^ at 3-d pONC, p<0.01). The 20% RGC reduction in the superior region was correlated with a 48% significant increase of axon bundle area in the same region (control: 94 ± 14.9 μm^2^, 3-d pONC: 140 ± 26.5 μm^2^, p=0.01). The 13% reduction in the nasal RGC density correlated with a 36% increase in the RGC axon bundle area (control: 91 ± 11.7 μm^2^, 3-d pONC: 124 ± 31.6 μm^2^), although the change was not significant (p=0.07, Fig. 7A). At 15-d pONC, axon bundles from both nasal (15-d pONC: 56 ± 15.7 μm^2^, p=0.2) and superior regions (15-d pONC: 54 ± 14.8 μm^2^, p=0.1) showed a reduction in area, which correlated with an 84% (nasal: 15-d pONC: 712 ± 52cells/mm^2^, p<0.001, student’s t-test) and 87% reduction in RGC density, respectively (superior: 15-d pONC: 414 ± 20 cells/mm^2^, p<0.001, student’s t-test, Fig. 7A).

**Figure 7.**
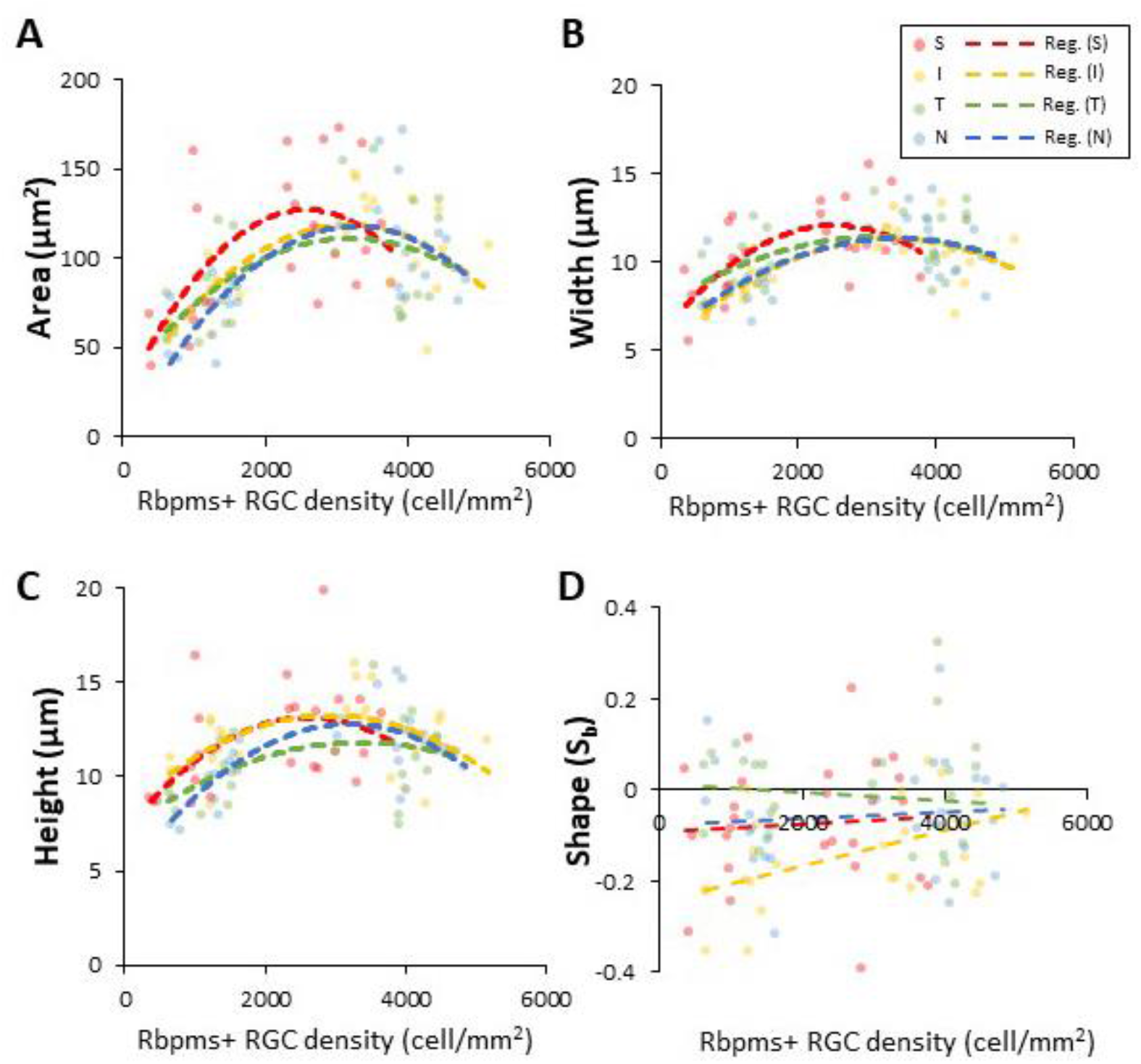
Regional differences were detected between RGC density and axon bundle size measurements. (A-D) RGC axon bundle cross-sectional area (A), width (B), thickness (C), and shape (D) plotted as a function of rbpms + RGCs density from each quadrant of each retina (S: superior; I: inferior; T: temporal; N: nasal). Second order polynomial regressions (dashed lines) were fitted to each region separately for (A-C), and linear regressions were fitted for (D).

**Table 2.**
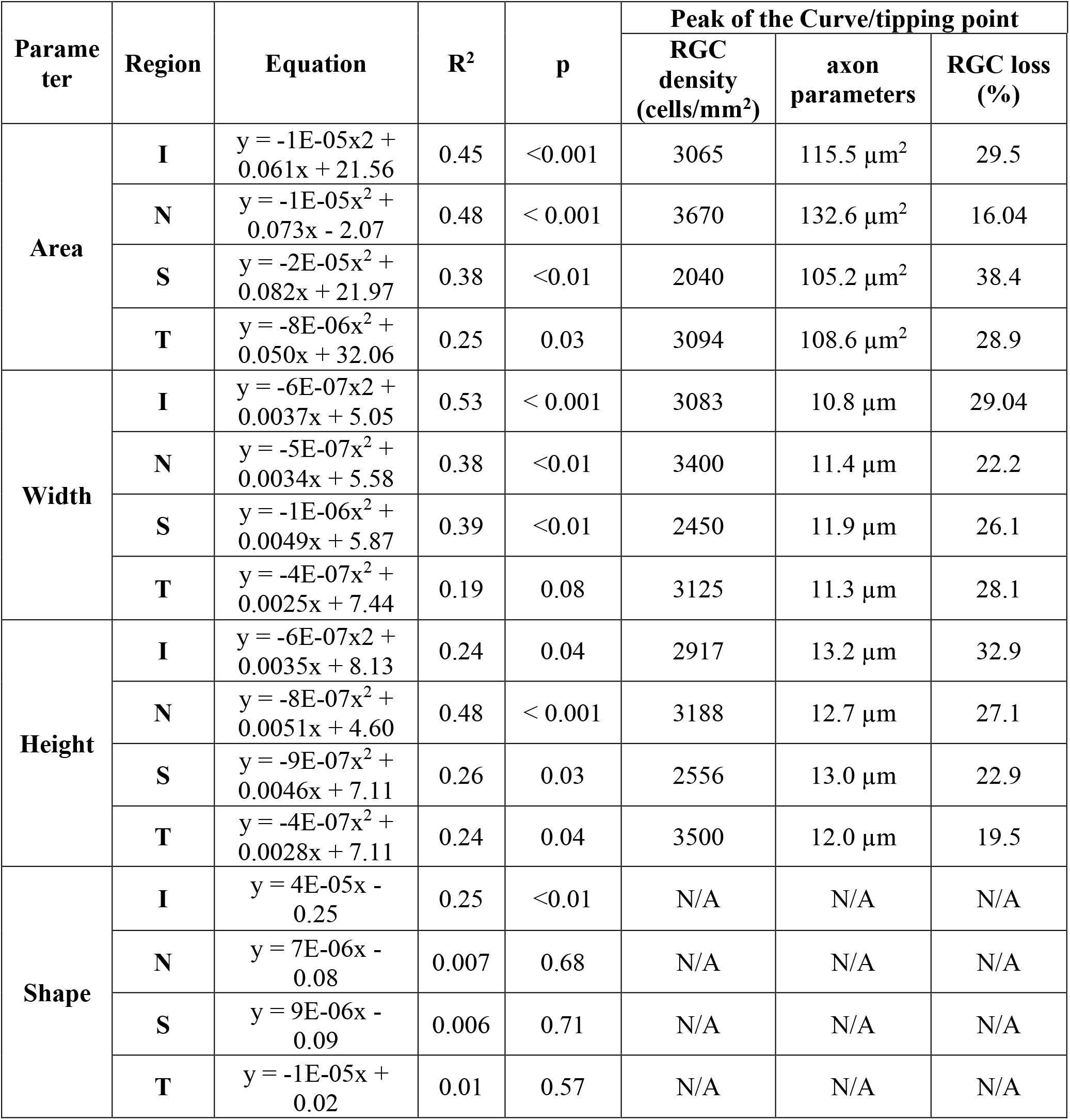
Region-dependent polynomial regression Analysis of RGC axon bundle morphology and soma loss (see Figure 6)

A linear regression model was used to fit the shape plot for each region (Fig. 7D). Although no regression was significant for superior, temporal, and nasal regions, a significant linear relationship was found between axon bundle shape and RGC density in the inferior region (R^2^ = 0.25, p<0.01), suggesting uneven changes in axon thickness and width following the ONC. Together, our data showed that the width, height, and area could serve as quantifiable biomarkers for RGC density globally and regionally.

### Numerical simulation suggests bundle area is most sensitive to RGC damage

We performed a numerical simulation to determine which bundle parameter quantified by vis-OCT is most sensitive to detect RGC damages. To model the change in RGC soma density as a function of days after ONC, we fit our experimental soma density data with a logistic function, as shown in Fig. 8A. Next, we simulated soma density values normally distributed along the modeled function within the recorded standard deviation. A total of 10 density values were simulated every 0.25 days, as shown in Fig. 8B. Simulated density values were used as input values for the bundle parameter models described in Fig. 6. Simulated parameter values were normally distributed along the model for each parameter using the experimentally recorded standard deviation values. The simulated parameter values were plotted as a function of time and an unpaired t-test (p>0.05) was used to determine at which time the parameter become significantly different from the baseline. Figures 8C-F show the change over time for simulated cross-sectional area (C), width (D), height (E), and shape (F). We repeated the simulation 50 times and the recorded average p-value as a function of time for each parameter. Figure 8G shows the simulation results for determining the most sensitive parameter (black dashed line indicates significance threshold): RGC density reaches the significance threshold at 1.3 days (blue), followed by bundle area at 9.6 days (purple), width at 10.1 days (orange), and height at 10.4 days (yellow). Unlike the other bundle parameters, bundle shape (green) never crosses the significance threshold, indicating that it does not have high enough sensitivity to detect a significant difference within 25 days of the ONC procedure.

**Figure 8.**
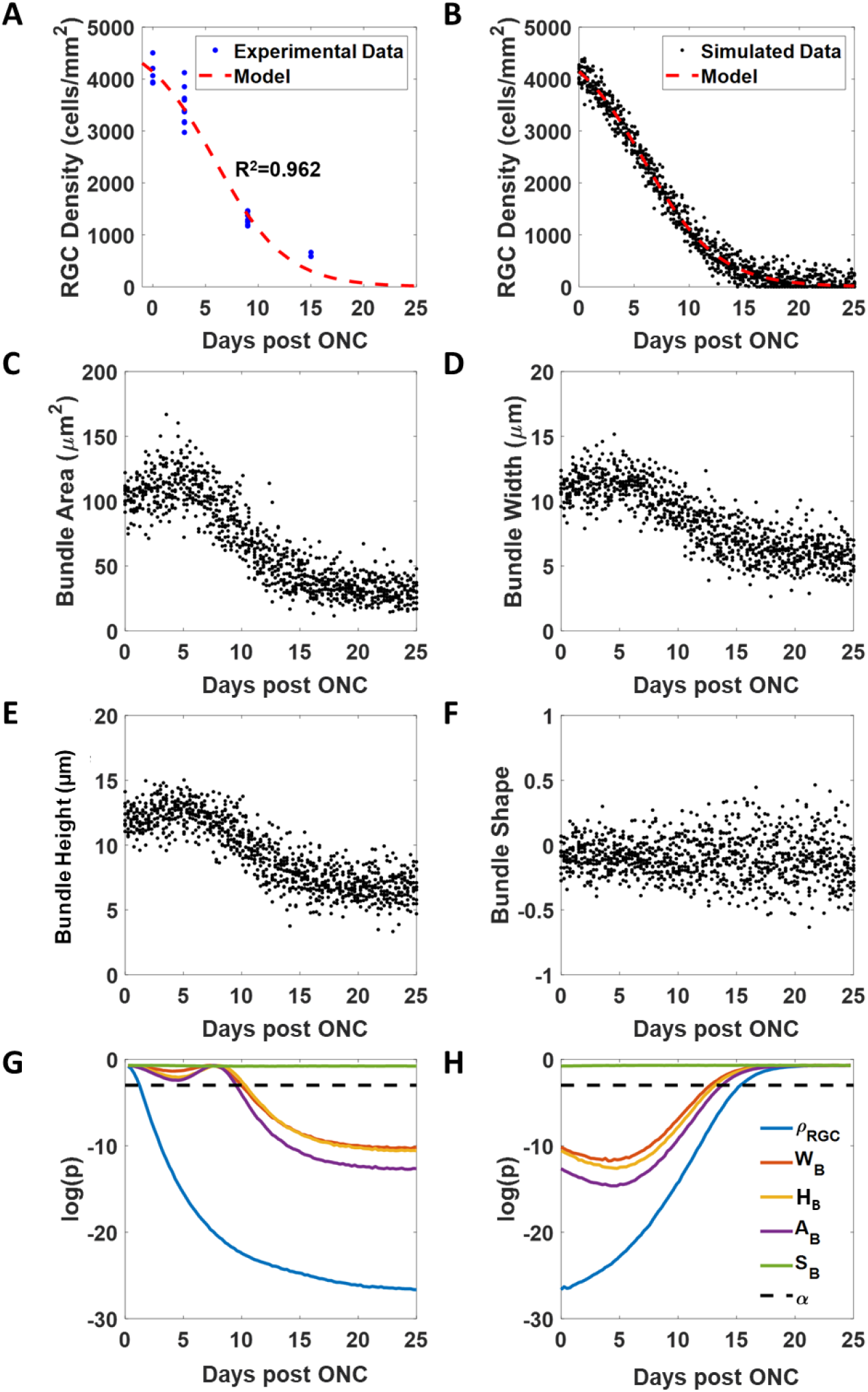
Simulation using experimental data to determine which RGC axon bundle size parameter is most sensitive to RGC damages and estimate parameter floor values. (A) RGC density (*ρ*_*RGC*_, blue data points) plotted as a function of time to establish relationship modeled by a logistic function (dashed red line). (B) Example simulated RGC density values (black data points) as a function of time. (C-F) Example simulated RGC axon bundle size parameters: cross-sectional area (A_B_) (C), width (W_B_) (D), thickness (T_B_) (E), and shape (S_B_) (F). (G) p-values as a function of time for the simulated size parameters to determine which parameters are most sensitive to RGC loss. (H) p-values as a function of time for the simulated size parameters to determine parameter floor times (*t*_*f*_).

The floor value is defined as the time point beyond which no further change in the respective parameter is detected [37, 38]. Using our simulated data, we estimated the floor value for each parameter. An unpaired t-test with a significance level of 0.05 was used to compare each simulated time point with the final time point. The time at which the p-value of the parameter crossed above the significance threshold was defined as floor time (*t*_*f*_). The mean of the parameter value beyond *t*_*f*_ was treated as the floor value. The simulation result for determining *t*_*f*_ for each parameter is shown in Fig. 8H: bundle width is the first to cross the significance threshold at 12.9 days (orange), followed by height at 13.4 days (yellow), area at 13.9 days (purple), and RGC density at 15.4 days (blue). The bundle shape parameter (green) did not cross the significance threshold and, thus, does not have a parameter floor value within 25 days of the ONC procedure. RGC density had a floor value of 131 cells/mm^2^, area had a floor value of 35.0 μm^2^, width had a floor value of 6.4 μm, and height had a floor value of 7.1 μm. The parameter floor model suggests that bundle area remains sensitive to size changes for the longest amount of time after crush because it reaches its *t*_*f*_ later than bundle width or height.

## Discussion

### RGC axon bundle morphology as a new *in vivo* biomarker for RGC damages

Glaucoma can progress without having easily identifiable symptoms until reaching an advanced stage of vision loss. Early diagnosis and intervention are crucial to slow down glaucoma progression [44]. Clinical OCT systems for the diagnosis and monitoring of optic neuropathies operate using NIR illumination. By shifting the illumination wavelengths to the visible light spectrum (510 nm to 610 nm), vis-OCT has an improved axial resolution of 1.3 μm in the retina compared to 4 μm in the best clinical NIR OCT devices [33]. In addition, vis-OCT has greater contrast between retinal layers due to the higher backscattering properties of biological tissues in the visible light wavelength range [33]. Taking advantage of this improved resolution and increased contrast sensitivity, we developed vis-OCTF to analyze individual RGC axon bundles *in vivo* [34-36, 40]. Thus far, we have demonstrated vis-OCTF in the mouse retina to visualize RGC axon bundles *in vivo* and validated these structures using confocal microscopy *ex vivo* [34]. We then applied vis-OCTF to visualize changes in RGC axon bundle structure in the case of increased RGC population using BAX^-/-^ mice and decreased RGC population using ONC mice [35]. In the present study, we developed new analytic tools for extracting RGC axon bundle size parameters from vis-OCTF images to determine which parameter is most sensitive to RGC damages.

Because every RGC extends one axon in the RNFL, we seek to examine whether directly quantifying changes of individual RGC axon bundles can be a more sensitive and accurate indicator than the bulk thickness of RNFL or GCIPL for RGC damage. In this study, we used an acute ONC injury model to longitudinally track morphological changes in single RGC axon bundles. We measured four bundle size parameters: (1) lateral width, (2) bundle height, (3) cross-sectional area, and (4) bundle shape. First, we found the bundle width from vis-OCTF is more sensitive to early damage compared to bundle height. The reduction of the lateral width was detected between 3-d and 6-d pONC (Figs. 3A - 3E), which was earlier compared to the reduction in the bundle height (6-d to 9-d pONC, Fig. 3B - 3E). Secondly, we introduced two novel parameters to measure and track the size and shape of single RGC axon bundles. By combining changes in both dimensions (lateral and axial), the cross-sectional area has shown to be an accurate indicator of RGC damage. At 3-d pONC, we found that 60% of all axon bundles showed a cross-sectional area increase of 30%, corresponding to about a 14% increase in width and a 15% increase in height (Figs. 3E-F). Thirdly, the bundle height measurements obtained following ONC matched the fact that RNFL thinning is not sensitive to the early neuropathic damage by ONC [22, 23]. We first observed a significant thinning of the RGC axon bundle at 9-days post-ONC (Figs. 3B-E). Importantly, we detected morphological changes in the RGC axon bundles but no significant change in the overall height of the GCIPL at 3-d pONC (Fig. 4), supporting the notion that RGC axon bundle morphology could serve as a more sensitive indicator than the GCIPL thickness.

RNFL swelling has also been reported [30], but *in vivo* changes in individual RGC axon bundles have not been demonstrated previously. Combining vis-OCTF with new analytic tools we have developed, we detected and quantified, for the first time, the swelling of individual axon bundles following acute ONC injury. Our results are in an agreement with the observation of a wide range of variations of changes in RNFL or GCIPL thickness right after disease insult. In addition, we found 15% of RGC soma loss correlated with the swelling in individual RGC axon bundles at 3-d post ONC. Interestingly, while axons were affected early in the disease, the rate of axon shrinkage was not as fast as that of the RGC somas. At 9-d pONC, we observed 68% of RGC soma loss while the axon bundles showed about a 12% decrease in cross-sectional area. At 15-d pONC, most RGC somas had degenerated (85%), while a substantial amount of axon bundles remained (30% decrease in axon bundle area). This desynchronization of soma and axon loss suggested a compartmentalized degeneration in RGCs, which also agrees with previous studies [45, 46].

### RGC and axon degeneration

Dendritic and axonal dysfunction is an early event in animal glaucoma models and may precede RGC soma degeneration [44, 47]. The ONH is hypothesized to be one of the most vulnerable structures to disease insult by glaucoma [44, 48, 49]. In our study, we observed obvious axon bundle swelling at 3-d pONC (Figs 2, 3, 5, and 6), an early time point when only 15% of RGC somas have degenerated. This suggests morphological changes in axon bundles occur early in disease progression, which agrees with previous studies [48]. In both DBA-2J mice, a genetic model of glaucoma [49], and a rat ocular hypertension model [48], swelling of individual axons was observed at early disease stages. Apart from individual axons, swelling of the RNFL [50] and retina [30] was also observed in early response to optic nerve injuries. While individual axon damage could lead to the swelling of the RNFL, other mechanisms may also contribute, including inflammatory responses such as macro and microglial proliferation in the RNFL [51-53].

We also show regional differences in bundle changes and RGC soma degeneration (Fig. 6). In WT retinas, the distribution of general RGCs varies across different regions [31, 54, 55]. For example, the density of rbpms positive RGCs in the superior region of the retina is 24% lower than the nasal retina in control mice. This difference in the overall RGC density could affect axon bundle organization and change how they react to disease insults. Furthermore, RGC types distribute unevenly across the retina [31, 54, 55], and different types of RGC respond differently to injury and disease [39, 55-57]. For example, one type of RGCs, the ipRGC, is involved in circadian functions [58], and at least one subtype, the M4 ipRGCs, has higher distribution in the superior and temporal retina [54]. It has been found that ipRGC’s are more resistant to chronic elevation of IOP [43] and ONC injury [59] than general RGCs. The retained circadian functions of the mice after chronic IOP elevation also suggest a normal function of ipRGCs. Therefore, it is likely that ipRGC axons suffered less damage and morphological changes than other RGCs in the case of ONC injury, potentially contributing to the regional difference in axon bundle morphology changes observed in our study. More studies remain needed to investigate whether the axons of RGC are organized into axon bundles based on function or location.

In summary, we established vis-OCTF parameters to track axon bundle morphology *in vivo* following the acute ONC injury. Our experimental and simulated results concluded that RGC axon bundle cross-sectional area is most sensitive to RGC damages. Our current study presented a new possibility to establish an *in vivo* quantifiable biomarker for RGC degeneration with glaucoma development and progression in future.

## Methods

Healthy adult (3-14 months) male and female wildtype (WT) C57BL/6 mice were used for this study. All animal protocols were approved by the University of Virginia institutional animal care and use committee and complied with the National Institutes of Health (NIH) guidelines.

### Optic nerve crush surgery

The ONC procedure was performed as described previously [35, 60]. Briefly, mice were anesthetized with an intraperitoneal injection of 100 mg/kg ketamine (Kataset, Zoetis; NADA #043-304) and 8 mg/kg xylazine (AnaSed, Akorn; NADA#139-236). A small incision was made in the superior and lateral conjunctiva, and the optic nerve was exposed by gentle dissection. The optic nerve was then gently clamped with a pair of forceps approximately 1 mm behind the globe for 7-10 seconds. After surgery, moxifloxacin (0.5%, NDC 60505-0582-4, Apotex Corp.) was applied to the crushed eyes to prevent infection. Mice were kept on a heating pad until fully recovered.

### *In vivo* vis-OCT imaging

A small-animal vis-OCT system (Halo 100; Opticent Health, Evanston, IL, USA) was used, as previously reported [34, 35]. In brief, four to six vis-OCT volumes (512 A-lines/B-scan × 512 B-scans/volume) were acquired from the same eye with the ONH aligned in each corner of the field of view (FOV) to cover different areas of the retina [35]. Each vis-OCT image volume was approximately 700 μm × 700 μm × 1500 μm (x × y × z). For each area, we generated vis-OCT fibergrams from the vis-OCT volume [34]. We used an intensity-based threshold method to detect the surface of the retina and cropped the RNFL by selecting the first ∼15 μm in depth, and then calculated the mean intensity projection along the axial (z) direction to generate the fibergram image composed of RGC axon bundles and surrounding vasculature. The fibergrams were then montaged, covering approximately 1.2 mm × 1.2 mm in total [35].

The vis-OCT volumes surrounding the ONH were digitally resampled to generate 425 μm radius circumpapillary B-scans. To do so, we manually marked the ONH in the enface image and plotted a ∼15 μm-thick arc around the ONH. The pixels were then sorted as a function of the angle measured between each sampled A-line and the nasal direction with the ONH as the vertex. Adjacent A-lines within a 0.1° sector were averaged together to reduce speckle noise while preserving spatial density.

### Individual axon bundle size quantification

We used the blood vessel pattern and the ONH as reference points to identify and track individual axon bundles in each retina (Suppl-Fig. 4). We measured the bundle height and lateral width of the individual RGC axon bundles using MATLAB. Bundle width measurements were recorded as previously reported. Briefly, the center axis of each bundle was manually marked, and the mean intensity profile along the center axis was plotted as shown in Fig. 1C. The intensity profile was normalized between 0 and 1, and the bundle width was recorded as the profile width at 1/e^2^. Bundle thickness was similarly recorded by extracting the axial intensity profile of the bundle and recording the thickness value at 1/e^2^, as shown in Fig. 1D. Measurement values were reported for individual bundles. Blood vessels were excluded from analysis by identifying the dark shadows in the B-scan images and uniquely distinguishable branching structures compared with surrounding axon bundles in fibergram images (Suppl-Fig. 4).

The elliptical approximation was used throughout this study instead of pixel-based measurements because of segmentation errors that occur when neighboring bundles are packed too closely together. To justify this approximation, we performed pixel-based measurements on isolated bundles and compared the results with the approximated area measurements. First, isolated bundles were manually selected from the resampled B-scans. Cropped images were contrast enhanced and median filtered to reduce noise. The filtered images were then binarized and morphologically opened to separate connected bundles. A watershed transform was applied and basins outside the selected bundle were removed. The pixel area of the segmented cross section was then calculated. Next, the cross-sectional shape of each bundle was approximated as an ellipse. Thus, the cross-sectional area of each bundle was determined as

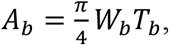

where A_b_ [μm^2^] is the bundle area; W_b_ is the bundle width [μm], and T_b_ is the bundle height [μm]. Finally, we compared the cross-sectional area measurements approximated by the area of an ellipse with pixel-base measurements of the same bundles and found no significant difference between the two methods (See Results).

We also developed a shape parameter to describe the width-to-height ratio of individual bundles using a dimensionless value normalized between -1 and +1. The shape parameter is defined as

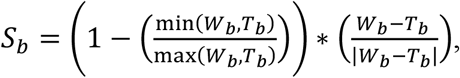

where S_b_ [dimensionless] is the shape parameter of a single RGC axon bundle. A positive S_b_ suggests a wider axon bundle elongated laterally, and a negative S_b_ suggests a taller bundle elongated axially.

To calculate the density of the axon bundles, we tracked the same area of each retina before and after ONC injury. We manually counted the number of bundles located at the radius of ∼425 μm in the fibergram images and calculated the bundle density per mm (Fig. 4). We also measured the GCIPL thickness, which was a measure of the top edge of the vitreous/RNFL and the bottom edge of the IPL, as indicated in Fig. 4A, blue arrows [35].

### Immunohistochemistry and confocal imaging

After acquiring vis-OCT images, mice were euthanized with 600 mg/kg euthasol (Euthasol, Virbac ANADA, # 200-071) and perfused with 4% paraformaldehyde (PFA) (ChemCruz, sc-281692). Eye cups were dissected, post-fixed in PFA for 30 minutes, washed with phosphate-buffered saline (PBS) containing Triton-X detergent (PBST, 0.5% Triton X-100), and then blocked for 1 hour in blocking buffer (1% BSA and 10% normal donkey serum, 0.5% Triton X-100 (Sigma-Aldrich, St. Louis, MO, USA). Primary antibodies, diluted using blocking buffer, included rabbit anti-rbpms (Abcam, ab194213, 1:500), mouse anti-neurofilament H antibody (Bio-Rad, MCA1321GA, 1:250), mouse anti-Tuj1 (gift from Tony Spano, University of Virginia, 1:250) and rat anti-Icam-II (BD Pharmingen, 553325, 1:500). Secondary antibodies, including donkey anti-mouse immunoglobulin G conjugated to Alexa Fluor 594 dye (Invitrogen A-21203, RRID: AB_141633) and donkey anti-rabbit immunoglobulin G conjugated to Alexa Fluor 488 dye (Invitrogen A-21206, RRID: AB_2535792), were diluted at 1:1000 in blocking buffer and incubated overnight at 4°C. After immunostaining, retinas were flat-mounted and cut into four quadrants: temporal (T), nasal (N), inferior (I), and superior (S). The blood vessel pattern was used as a landmark to align vis-OCTF and confocal images of flat-mounted retinas (Suppl-Fig. 4).

Confocal images were taken using a Zeiss LSM800 confocal microscope (Zeiss, Thornwood, NY). Z-stack images covering the depth of the outer nuclear layer (ONL) to the GCL (approximately 50-80 μm) were acquired. Lower magnification (5×) pictures were captured for the whole retina using the tiling/stitch function in Zen (Zen 3.2; Oberkochen, Germany). For cell counting, individual images were captured at 10×, covering an area of 0.408 mm^2^. To show morphological changes in degenerating axons, individual z-stack images covering an area of 0.0037 mm^2^ were taken using a 63× water immersion objective.

### RGC soma quantification

We followed our published protocols to quantify RGC soma density [39, 43, 61, 62]. In brief, the nasal side of the eye was marked during eye dissection for orientation. To quantify RGC density, mouse retinas were immunostained with anti-rbpms antibody. For each retina, 16 enface z-stack images covering the depth of GCL were captured (20-30 μm of total depth). Four images were acquired for each of the four quadrants to ensure broad coverage of the entire retina (Fig. 5A). Three rectangles covering no less than 0.03mm^2^ area were randomly drawn on the images, avoiding overlaps with large blood vessels. All rbpms-positive cells within the rectangles were manually counted in Zen (Zen 3.2; Oberkochen, Germany) [43]. Cell density was calculated using total cell counts divided by area for the four quadrants in each retina. All data analyses were cross-examined by two independent observers.

### Statistical analyses

All statistical analyses were performed using MATLAB and Prism. We used a linear mixed-effects model for all height, width, and area comparisons to remove influence from individual subjects. Two sets of ONC experiments were performed. One set was to track changes in the axon bundle parameters following the ONC of the same retina. The other set was to correlate changes in axon bundle morphology with RGC soma loss using different retinas. For both sets of experiments, we performed one-way analysis of variance (ANOVA) followed by Dunnett test for multiple comparisons. A significance level of 0.05 was used and the p-values of the Dunnett tests were reported unless otherwise stated. All results were reported as mean ± standard deviation.

## Acknowledgments

This work was supported in part by NIH grants R01EY029121, R01EY019949, U01EY033001, and R44EY026466. We thank William Tucker for his technical support on bundle quantification and Prof. Ignacio Provencio for his insightful discussions.

## Supplementary Figures

**Suppl-Figure 1.**
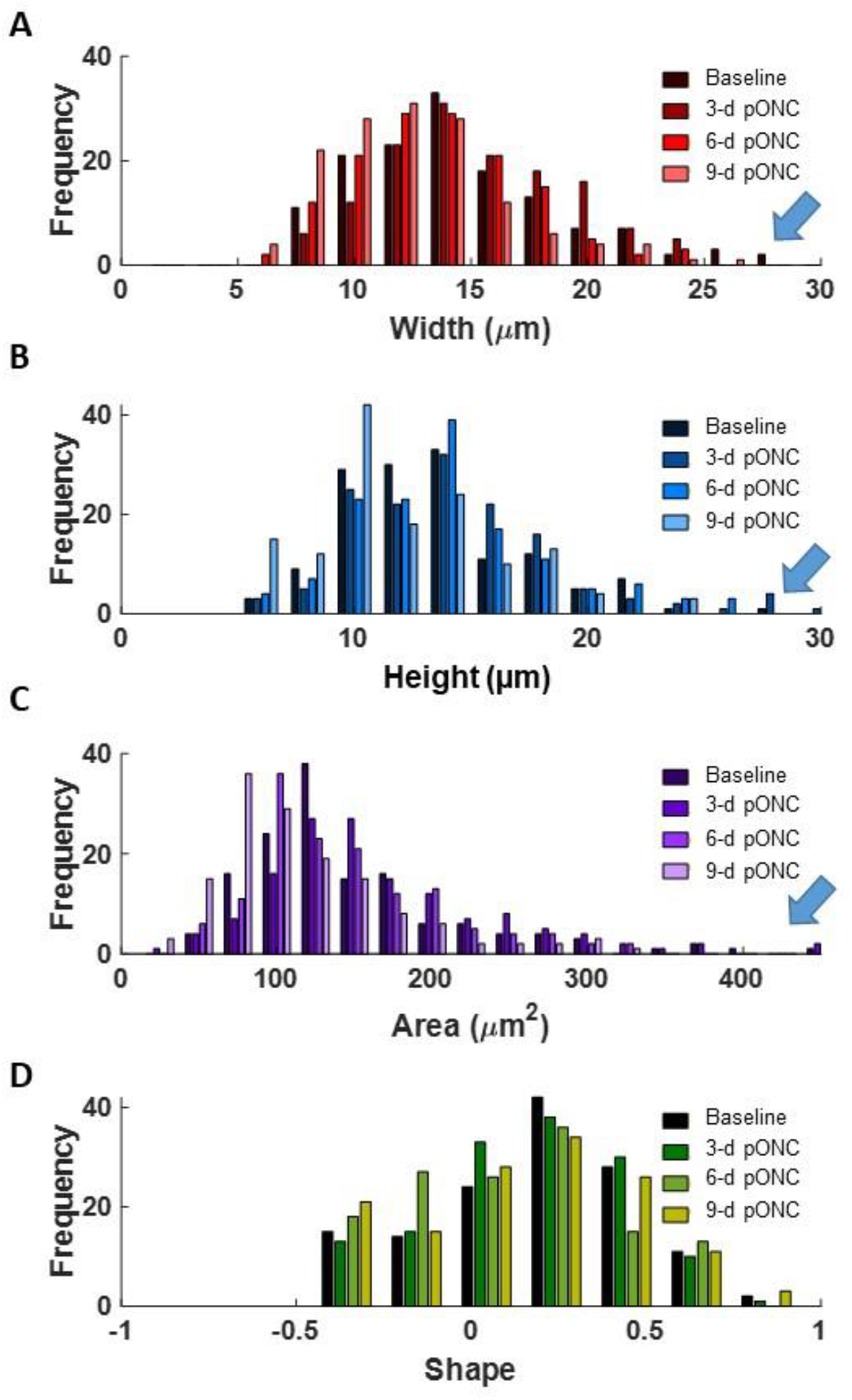
Histogram of tracking of a total of 141 RGC axon bundles before ONC and 3-d, 6-d, 9-d pONC (See Figure 3). The width (A), height (B), area (C), and shape (D) were measured, and values were plotted as shown in the histograms. Blue arrows indicate large axon bundles that decrease in size with time after ONC.

**Suppl-Figure 2.**
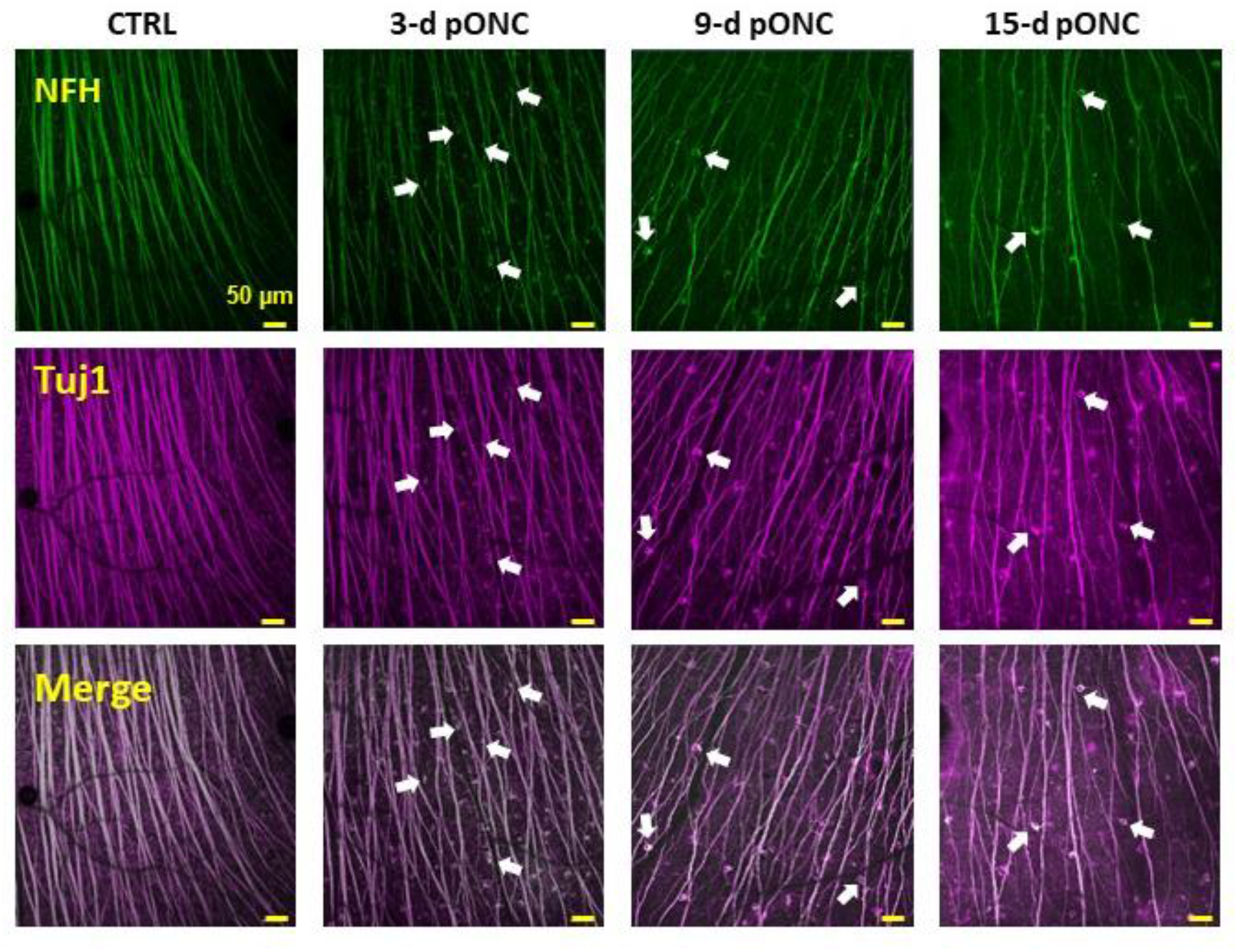
Degenerating RGCs were observed post ONC. Flat mounted retinas were double immunostained with Tuj-1 (purple) and NFH (green, see Figures 7 and 8). Confocal images were taken in the medial to peripheral area of the retina. Tuj-1 labeled axon bundles and some RGC soma membranes, while NFH antibody labeled axon bundles and degenerating RGC somas (white arrows).

**Suppl-Figure 3.**
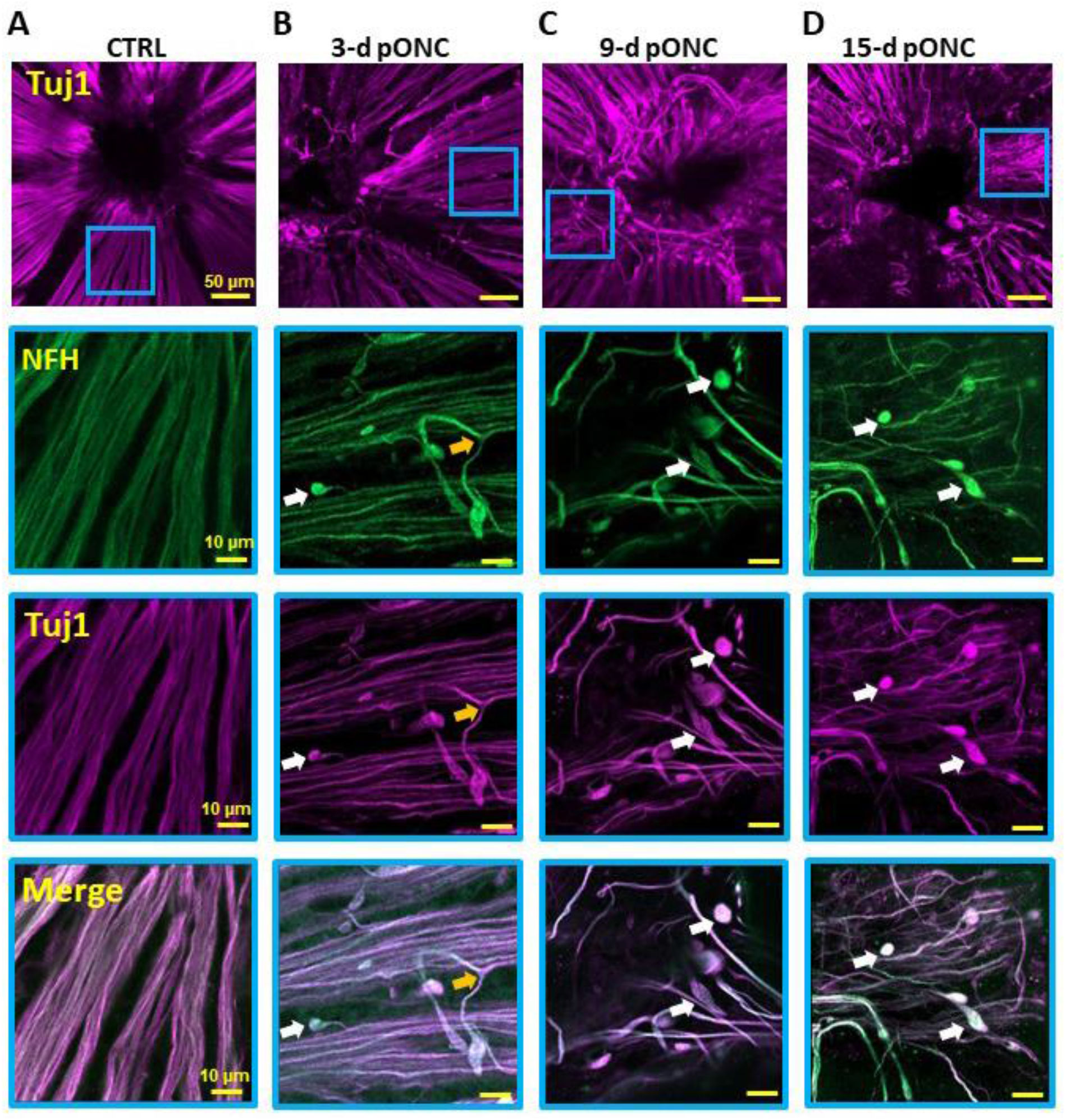
Confocal microscopy images of RGC axon bundle morphological changes after ONC injury. Images of flat mounted retinas of control (A), 3-d (B), 9-d (C) and 15-d (D) pONC were double-immunostained with Tuj-1 (purple) and NFH (green) antibodies. Low magnification images taken near the ONH (top panels). High magnification airy scan microscopy images acquired at areas indicated by blue rectangles immunostained by NFH (middle panels) and Tuj1 (bottom panels). The yellow arrow indicates a splitting axon bundle, and the white arrows indicates retraction bulbs that formed at axon lesion sites.

**Suppl-Figure 4.**
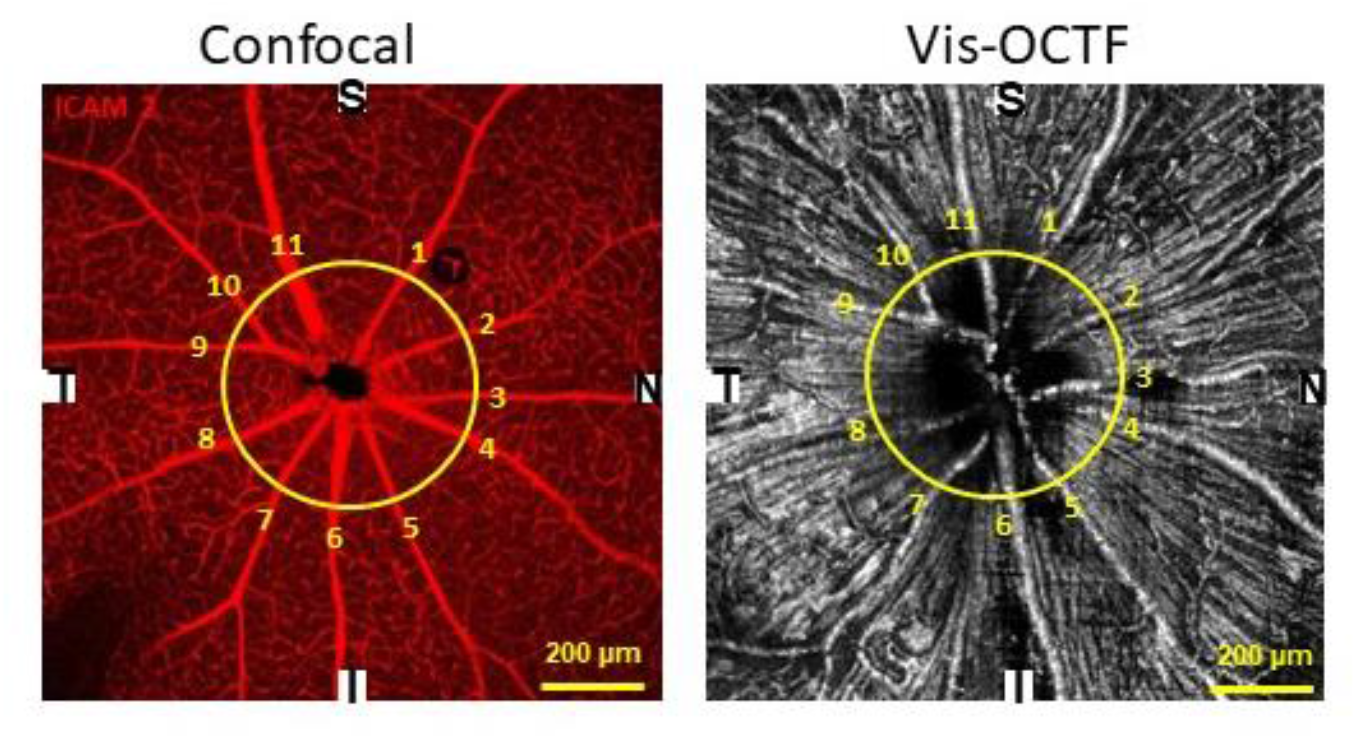
Example of how blood vessel pattern was used as a landmark to align vis-OCTF and confocal images of flat mounted retinas (See Figure 5). The same blood vessels were labeled (1-11) in the *ex vivo* confocal image of flat-mounted retina immunostained for ICAM2 and *in-vivo* vis-OCT fibergram of the same retina. S: superior; N: nasal; T: temporal; I: inferior.

